# Autoreactive antibody production by intrarenal B cells in mouse kidney allograft rejection

**DOI:** 10.64898/2026.06.15.732429

**Authors:** Ismail Sayin, Jong Cheol Jeong, Deepjyoti Ghosh, Samarth S. Durgam, Jacqueline BM Oien, Alexander J. Nelson, Dengping Yin, Peter T. Sage, Anat R. Tambur, Marcus R. Clark, Madeleine S. Torcasso, Anita S. Chong

## Abstract

Antibody-mediated rejection (AMR) is a major cause of kidney allograft failure, with donor HLA-specific antibodies (DSA) recognized as primary drivers of AMR pathology. However, a significant proportion of AMR diagnoses lack detectable DSA, thus implicating DSA-independent mechanisms. In support, we previously reported on the accumulation of autoreactive B cells in rejecting renal biopsies, which raised the possibility that autoreactive IgG produced within the allograft contributes to graft pathology. In this study, we used mouse models to show that rejecting kidney allografts preferentially promoted a breach in autoreactive B cell tolerance, leading to the local production of autoreactive antibodies. Notably, autoreactive IgG responses were uncoupled from DSA production, indicative of distinct regulation. Organoid cultures confirmed that autoantibody production was observed in the rejecting kidney, whereas DSAs were produced in both the lymph node and graft. Intrarenal B cells expressing Nur77 were enriched for autoreactivity, consistent with *in situ* antigen recognition. Finally, autoreactive antibodies are pathogenic, since inhibiting autoantibody production with transient anti-IL-15 and CTLA-4Ig led to preserved kidney allografts. These unique features of *in situ* autoantibody responses may be relevant to diverse diseases with chronic tissue inflammation beyond transplantation.

## Introduction

Antibody-mediated rejection (AMR) is the leading cause of immune-mediated late allograft failure in kidney transplantation, where the detection of circulating donor-specific antibodies (DSA) together with distinguishing histologic and molecular features are the current diagnostic criteria for AMR^1,2^. Nevertheless, a substantial fraction of kidney allograft biopsies with molecular and histological features of AMR lack detectable circulating DSA, thus raising the possibility that additional antibody specificities, including natural, non-HLA-specific and/or autoreactive antibodies may mediate AMR ^3–7^. Innate immune mechanisms that do not involve antibodies have also been implicated ^8–10^. However, recent transcriptional analysis of AMR biopsies with or without circulating DSA indicate broad transcriptional overlap^1,11,12^. This suggests that non-HLA antibodies or HLA antibodies produced locally within the allograft are the primary drivers of AMR when circulating DSA is not detected.

One approach to probe the mechanisms driving DSA-negative AMR is by interrogating the specificity of B cells infiltrating human renal allograft biopsies. Using single cell RNA-sequencing approaches to probe B cells infiltrating biopsies diagnosed with chronic or chronic and active AMR, Asano et al. reported that they expressed an innate cell transcriptional state resembling mouse peritoneal B1 or B-innate cells ^13^. Notably, antibodies produced by the intrarenal B cells did not bind to donor HLA or to ubiquitously expressed self-antigens; instead, they were reactive to renal-specific or inflammation-associated antigens ^13^. Similar findings of intra-graft accumulation of innate B cells and autoantibodies were also reported in lung transplant rejection ^14,15^. Furthermore, in murine kidney transplantation, less than half of intrarenal class-switched B cells are donor-specific, suggesting the remainder may be autoreactive^16,17^. Together, those observations suggest that local antigens within the inflamed kidney allograft provide signals that overcome B-cell tolerance, prompting the proliferation of intra-graft B cell and differentiation into plasma cells. The mechanisms leading to the loss of autoreactive B cell tolerance in individuals with autoimmune disease are characterized by defects in central or peripheral tolerance mechanisms ^18,19^, but whether these mechanisms also mediate the loss of autoreactive B cell tolerance in transplant recipients has not been characterized.

Autoreactive B cells detected in healthy mice and humans are enriched for polyreactivity to apoptotic cells and/or commensal bacteria ^20,21,22^. Furthermore, checkpoint inhibitors such as anti-PD-1 and anti-CTLA4 used to treat cancers have also been reported to induce autoimmune-related adverse events and permit the emergence of autoreactive mature B cells ^23^. In addition, an increasing number of studies have reported on polyreactive and autoreactive antibodies correlating with worse graft outcomes in solid organ transplant rejection ^6,24–26^. These autoantibodies recognize Angiotensin II Type 1 Receptor (AT1R), tubulin, collagens, vimentin, tissue-restricted or endothelial cell targets, and have been associated with microvascular injury and chronic AMR rejection ^26–29^. Nevertheless, multiple gaps in knowledge persist with regards to the immunobiology of non-HLA antibodies in transplant rejection. These include when and where autoantibodies are produced, whether autoreactive B cells undergo epitope spreading so that their antigen specificity evolves over time post-transplant, and if antigen specificity determines autoantibody function and resulting graft pathology.

In this study, we used mouse models of kidney allograft rejection to address some of these gaps in knowledge. We show that autoantibodies were produced independently of DSA and within rejecting kidney allografts. Furthermore, while transient CTLA-4Ig prevented the production of DSA long-term it did not prevent autoantibody production. In contrast, the combination CTLA-4Ig with anti-IL-15 was able to prevent autoantibody production and preserve kidney allograft pathology.

## Results

### Transient CTLA-4Ig suppresses donor-specific but not autoreactive IgG responses

We first quantified DSA and autoantibody production in C57BL/6 recipients of BALB/c kidney allografts undergoing unmodified rejection, or in recipients treated with transient CTLA-4Ig (on day 0 and 7). Donor-specific IgG was quantified using BALB/c splenocytes (Fig. 1A) or donor MHC class I and class II–coated multiplex beads (Fig. 1B-C, ^30^). Robust anti-BALB/c IgG responses were observed in untreated recipients, with maximum anti–class I and anti–class II DSA detected by 2 weeks post-transplant (Fig. 1A–C, Fig. S1A–E). In contrast, recipients treated with transient CTLA-4Ig displayed markedly reduced DSA for up to 8 weeks post-transplant (Fig. 1A–C, Fig. S1). IgG responses to anti-donor Class I were most effectively inhibited, whereas a modest increase in donor class II, I-E^d^, was detected in these recipients of transient CTLA-4Ig.

**Figure 1.**
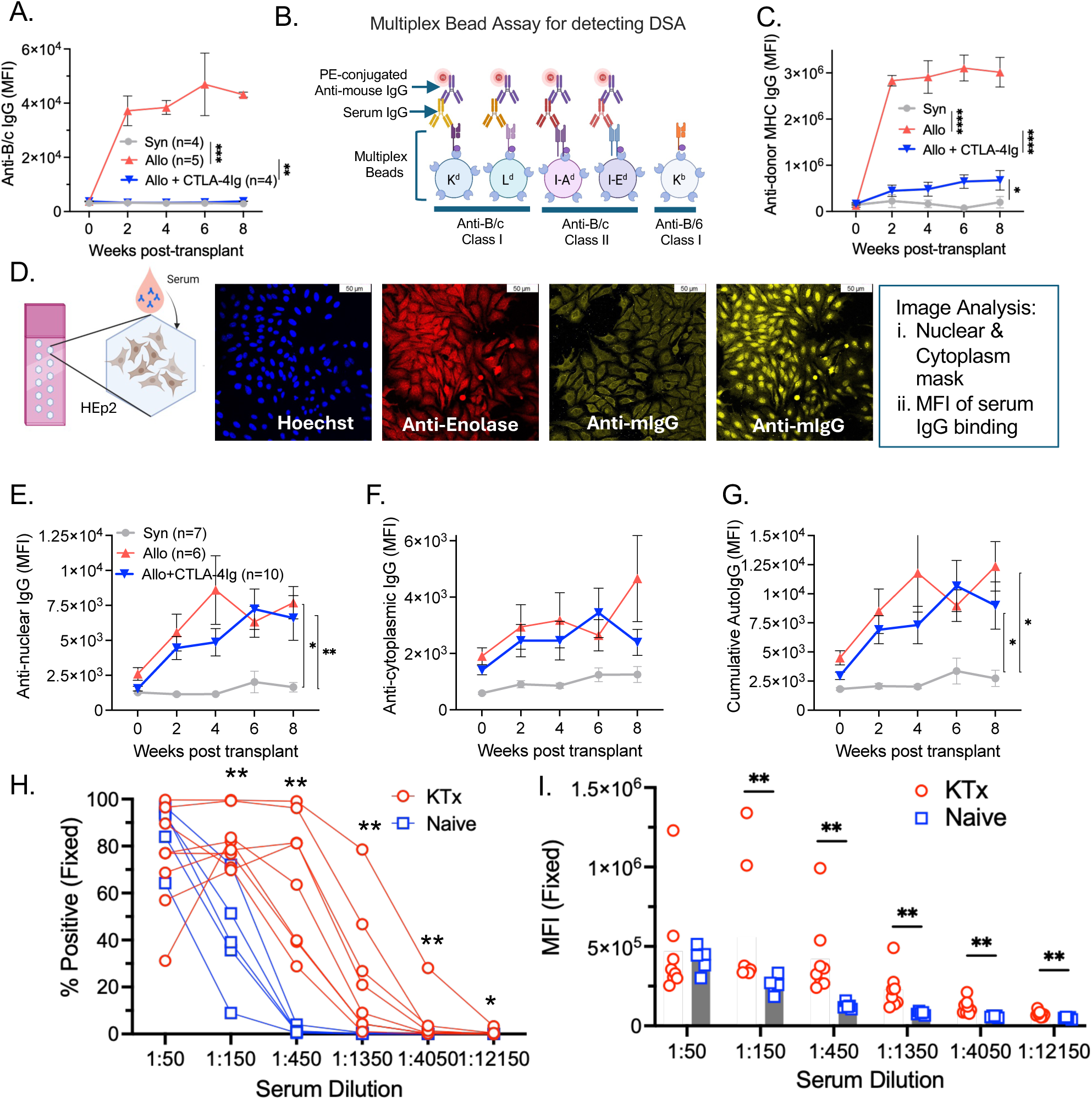
Transient CTLA-4Ig suppresses donor-specific but permits autoreactive IgG responses. C57BL/6 mice received kidney transplants from C57BL/6 (Syn) or BALB/c (Allo) donors. Some BALB/c kidney recipients received CTLA-4Ig (250 µg/mouse on D0 & 7). (A) Levels of serum anti-BALB/c IgG quantified by flow cytometry using BALB/c splenocytes as targets. Data are presented as mean fluorescence intensity (MFI) (n = 4 for Syn or Allo + CTLA-4Ig, n = 5 for Allo). (B) Multiplex bead assay for detecting anti-donor MHC Class I (K^d^, L^d^) and Class II (I-A^d^, I-E^d^) (DSA) by flow cytometry. (C) Levels of serum cumulative DSA. Data presented as MFI (n = 4 for Syn & Allo + CTLA-4Ig, n = 5 for Full mismatch). (D) Schematic for quantifying serum autoreactive IgG using Hep-2 ANA kit. (E-G) MFI of autoreactive IgG against nuclear (E), cytoplasmic (F) and cumulative nuclear + cytoplasmic antigens (G). n = 6 for Syn or Allo, n = 10 for Allo + CTLA-4Ig) (H-I) Flow cytometric assessment of autoreactive IgG to fixed HEp2 cells. Sera from naïve or Allo + CTLA-4Ig treated kidney transplanted mice (KTx) were serially diluted in PBS, and data are presented as (H) % positive or (I) MFI. Data are presented as mean ± SEM, and statistical significance was assessed by Mann-Whitney and Mixed-effects analysis. *P < 0.05; **P < 0.01; ***P < 0.001.

Autoreactive IgG was quantified using fixed and permeabilized HEp-2 cells from an commercial anti-nuclear antibody (ANA) kit, followed by image analysis to quantify IgG binding to nuclear and/or cytoplasmic antigens (Fig. 1D, ^13^). Untreated recipients produced both anti-nuclear and anti-cytoplasmic IgG, with peak anti-nuclear IgG observed at 4 weeks post-transplant (Fig. 1E-G). Strikingly, despite effective inhibition of DSA, CTLA-4Ig–treated recipients developed robust HEp-2–reactive IgG comparable to untreated recipients, with peak anti-nuclear and anti-cytoplasmic responses observed at 6 weeks post-transplant (Fig. 1D–G). These findings indicate that autoantibody responses can occur independently, and with slightly delayed kinetics compared to DSA responses. To confirm that autoantibody production can be independent of DSA responses, we examined autoantibody production in C57BL/6 recipients of kidneys from bm12 mice that differ at only three nucleotide differences in the I-A beta genes^31^. As previously reported^32^, this model results in minimal or no anti-bm12 IgG but substantial autoreactive IgG responses (Fig. S2).

Immunofluorescence followed by image analysis is labor intensive and has a relatively narrow dynamic range, so we developed a more tractable and quantitative flow cytometry-based assay using fixed and permeabilized HEp-2 cells. With this assay, we conducted serial dilutions to compare the titer of autoreactive antibodies in sera from naive mice or kidney transplant recipients and distinguish autoreactive antibodies in transplant recipients from low-affinity IgG in naive sera. We confirmed that sera from kidney transplant recipients had significantly increased titers of autoreactive IgG compared to naïve sera, based either on percentage of positive cells or mean fluorescence intensity (MFI) (Fig. 1H, I, Fig. S3). Together, these imaging-based and flow cytometric assays demonstrate that autoreactive IgG production can be uncoupled from DSA during allograft rejection.

### HEp-2–reactive IgG recognizes autoantigens in kidney tissues

To address potential concerns arising from the use of human epithelial HEp-2 cells, we tested if serum samples that bound to HEp-2 cells were also reactive to mouse kidney tissues (Fig. 2A). To minimize background from endogenous IgG binding to mouse tissues, we used C57BL/6 kidneys from *CD19*^Cre/+^*Prdm1*^flox/flox^ mice that have defects in plasma cell differentiation and thus, minimal circulating Ig ^33^. Tissue sections from healthy kidneys and kidneys that had been subjected to ischemia–reperfusion injury (IRI) were used. Immunohistochemistry was performed to detect autoreactive IgG in sera from naive mice, as well as untreated and CTLA-4-treated kidney transplant recipients (n=26 serum samples that bound to HEp-2 cells with a range of MFIs). Robust IgG binding was observed to both nuclear and cytoplasm regions of cells within the renal cortex of naive and IRI kidney sections (Fig. 2B). Quantitative image analysis of IgG binding to kidney tissue showed that sera samples (n=4-5 of 26) that weakly bound to HEp-2 (MFI<7000) also weakly bound to normal or IRI kidney (MFI<1800). These data indicate that the HEp-2 assay yielded minimum/no false positives (Fig. 2C,D). We also identified a fraction of samples that weakly bound to HEp-2 but strongly bound to kidney sections (n=7-8 of 26), suggesting that these sera samples were recognizing antigens present only in mouse kidney cells. Finally, of the 13 samples that bound to both HEp-2 cells and kidney tissues, we observed a significant correlation in MFI consistent with shared autoantigens expressed by mouse kidney and HEp-2 cells (Fig. S4A,B). Together, these data show that autoreactive IgG detected in the HEp-2 assay recognize autoantigens expressed by mouse kidney cells, but that approximately 30% of sera that were autoreactive to mouse kidney sections were missed by the HEp-2 assay.

**Figure 2.**
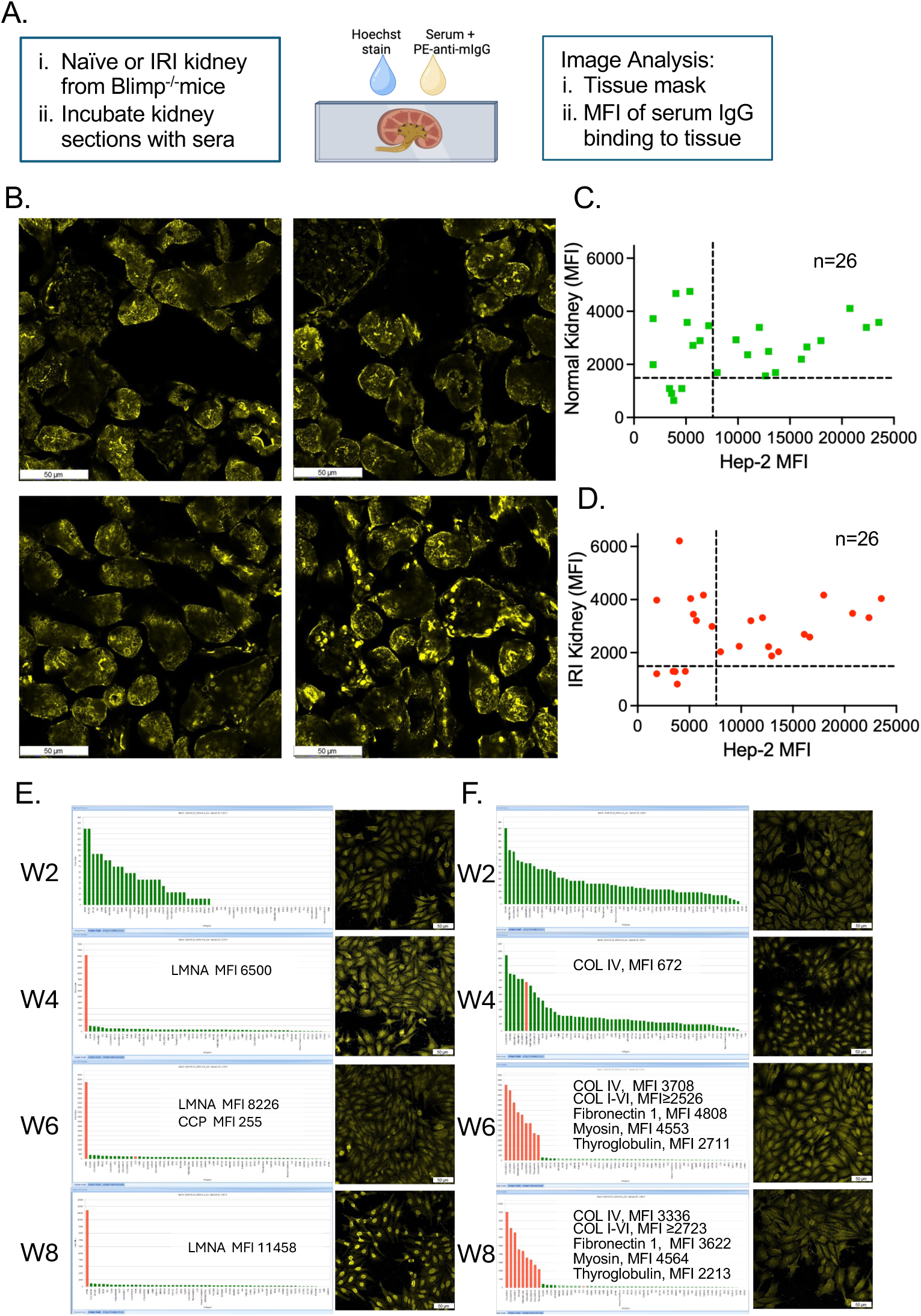
Hep-2-reactive IgG recognizes autoantigens in kidney tissues. (A) Schematic of autoantibody detection on syngeneic mouse tissue. Sera from untreated and CTLA-4Ig-treated recipients collected at week 0-8 post-transplantation, and that bound to HEp-2 cells with a range of MFIs. (B) Representative confocal images of autoreactive IgG bound to normal (top) or IRI (bottom) kidney sections. (C-D) Correlation of syngeneic tissue binding and Hep-2 ANA detection kit binding on native and IRI-induced kidney sections (n=26 matching serum samples). (E-F) Serial serum samples from two BALB/c kidney transplant recipients that bound to LifeCode non-HLA beads (left panels) and HEp-2 cells (right panels). Red bars indicate positive binding to beads.

Commercial multiplex bead assays are being used to quantify non-HLA antibodies in clinical transplantation ^34–36^. We tested whether the LifeCode non-HLA bead assay, which displays 60 autoantigens, was able to identify sera that had been identified as positive for autoreactive IgG in the HEp-2 assay. We observed that most of HEp-2-positive sera from CTLA-4Ig-treated kidney recipients did not bind any of the non-HLA beads consistent with HEp-2 cells providing a much broader range of antigens than the bead assay (Table 1). Sera from one recipient bound to Lamin A at week 4 post-transplant, and with increasing MFI over time post-transplant (Fig. 2E). Sera from another recipient demonstrated epitope spreading, starting with reactivity to Collagen IV on week 4 post-transplant, and reactivity spreading to Collagen 1-VI, fibronectin, myosin and thyroglobulin and with increasing MFI over time post-transplant. (Fig. 2F). These data demonstrate that HEp-2 assay captures a broader repertoire of non-HLA autoantibodies, while the bead assay better reveals the evolution of the autoreactive antibody repertoire post-transplant, notwithstanding the caveat that the bead assay displays human antigens, as do HEp-2 cells.

**Table 1.** Detection of autoantibodies with HEp2 cell immunofluorescence assay versus Lifecode Non-HLA autoantigen kit beads. Serum from 5 kidney transplant recipients treated with CTLA-4Ig (Allo+CTLA-4Ig) and harvested on the indicated days post-kidney transplant were tested on HEp2 cells and Lifecode beads. C: Cytoplasmic; N: Nuclear binding on HEp2 cells, and Negative or Positive indicate staining on HEp2 or Lifecode beads.

| Mouse ID | Week Post-KTx | HEp2 | Lifecode |
| --- | --- | --- | --- |
| 1189 | 2 | C+N | Negative |
| 1189 | 4 | N | Positive |
| 1189 | 6 | C+N | Positive |
| 1189 | 8 | Negative | Positive |
| 1197 | 2 | C+N | Negative |
| 1197 | 4 | C+N | Positive |
| 1197 | 6 | C+N | Positive |
| 1197 | 8 | C+N | Positive |
| 2898 | 2 | C+N | Negative |
| 2898 | 4 | C+N | Negative |
| 2898 | 6 | C+N | Negative |
| 2898 | 8 | N | Negative |
| 205 | 2 | N | Negative |
| 205 | 4 | N | Negative |
| 205 | 6 | Negative | Negative |
| 205 | 8 | Negative | Negative |
| X | 2 | C+N | Negative |
| X | 4 | C+N | Negative |
| X | 6 | C+N | Negative |
| X | 8 | N | Negative |

### Rejecting kidney grafts are the site of autoantibody production

Observations of Asano et al. ^13^ that autoreactive B cells preferentially accumulate in rejecting kidney biopsies, and that in murine kidney transplantation, less than half of intrarenal class-switched B cells are donor-specific, suggest the remainder may be autoreactive^16,17^, and that autoantibodies may be preferentially produced in rejecting kidney grafts. To test this, we generated tissue fragments from the cortical region of transplanted kidney explants or from draining lymph nodes retrieved from untreated or CTLA-4Ig–treated recipients, as previously described ^37^. After a 4-day culture, donor-specific and autoreactive IgG in the organoid supernatants were quantified (Fig. 3A). Culture supernatants of lymph nodes from untreated recipients at week 2 or 7 post-transplant had significantly increased donor-specific IgG compared to lymph nodes from naive mice, consistent with DSA originating from the lymph node. Consistent with the lack of serum DSA, culture supernatants of lymph nodes from CTLA-4Ig-treated recipients had minimal DSA (Fig. 3B). In contrast, autoreactive IgG was not significantly increased in the majority of supernatants from lymph nodes of untreated or CTLA-4Ig treated recipients, with the exception of a few lymph node cultures from untreated recipients (Fig. 3C).

**Figure 3.**
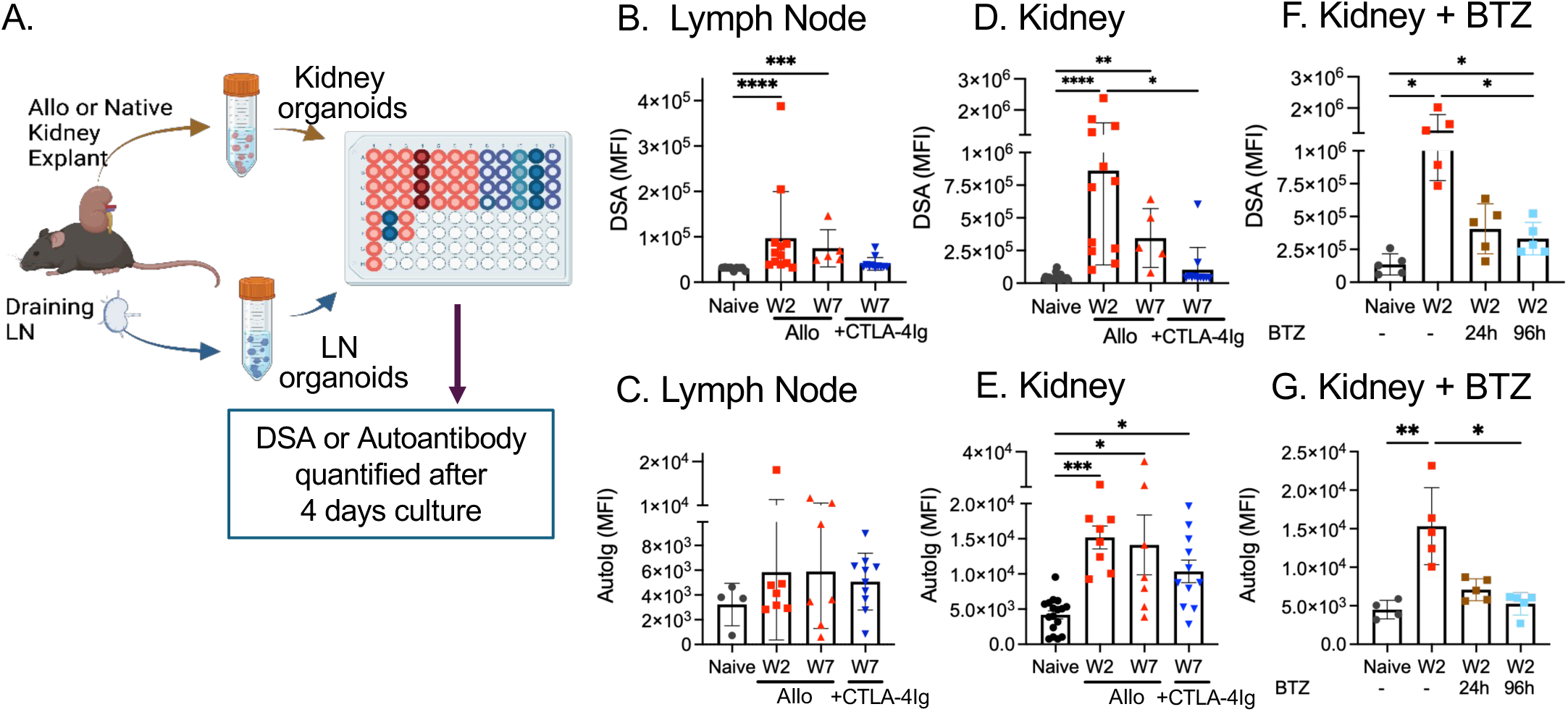
Rejecting kidney grafts are the site of autoantibody production. (A) Schematic of DSA and autoantibody detection in the culture supernatants of draining lymph nodes (LN) or kidney organoids from untreated (Allo) or CTLA-4Ig-treated (+CTLA-4Ig) recipients of BALB/c kidneys. (B-D) Anti-donor MHC IgG (DSA) measured by flow cytometry with multiplex MHC-coated beads. (E-G) Autoreactive IgG (AutoIg) measured with HEp-2 ANA detection kit. (D, G). Kidney organoids from W2 allograft recipients were cultured with proteosome inhibitor, bortezomib (BTZ), added at the beginning of the culture for 24h or 96h to deplete antibody-secreting cells. Data are presented as mean MFI ± SEM. Statistical significance was assessed by ordinary ANOVA test. *P < 0.05; **P < 0.01; ***P < 0.001; ***P < 0.0001.

Kidney organoids from untreated recipients harvested at week 2 post-transplant produced DSA, and this production was significantly reduced by week 7 (Fig.3D). As expected, CTLA-4Ig treated recipients produced little or no DSA. In contrast, autoreactive IgG was detected in kidney organoid cultures from all experimental groups including from CTLA-4Ig treated recipients (Fig. 3E). To confirm that the IgG in the supernatants were produced by antibody-secreting cells within the kidney organoids, we added the proteasome inhibitor bortezomib at the start of the culture to deplete antibody-secreting cells (Fig. 3F-G ^38,39^). When bortezomib was added for the first 24 h or duration of culture, the amount of DSA or autoreactive IgG was significantly reduced, confirming that antibodies were produced *in situ* by antibody-secreting cells. Together, these data show that the rejecting kidney allograft harbors antibody-secreting cells that locally produce donor-specific and autoantibodies, whereas DSA is produced in both the lymph node and graft

To characterize the cellular context of intrarenal antibody production, we analyzed the immune cell infiltrates in rejecting kidney grafts. Flow cytometric analysis of the immune infiltrate revealed a significant increase in CD19^+^ B cells in CTLA-4Ig–treated recipients compared to native kidneys (Fig. 4A; Fig. S5). Among intrarenal B cells, we observed an enrichment of germinal center (GC)-like B cells (GL7^+^CD95^+^) ^40–42^, especially within the donor-MHC-negative B cell population, suggesting that the majority of GC-like B cells were not donor MHC–specific (Fig. 4B–E). Intrarenal CD4^+^ T cells showed an increased frequency of CXCR5^+^PD-1^+^ Tfh-like cells in rejecting grafts relative to controls (Fig. 4F ^16,43^). Consistent with these findings, immunofluorescence imaging of rejecting kidney grafts revealed accumulation of B cells and CD4⁺ T cells into aggregates within the kidney cortex parenchyma and frequently localized in perivascular regions (Fig. 4G). These data suggest that local GC-like interactions within the graft may contribute to the *in situ* generation of autoreactive IgG.

**Figure 4.**
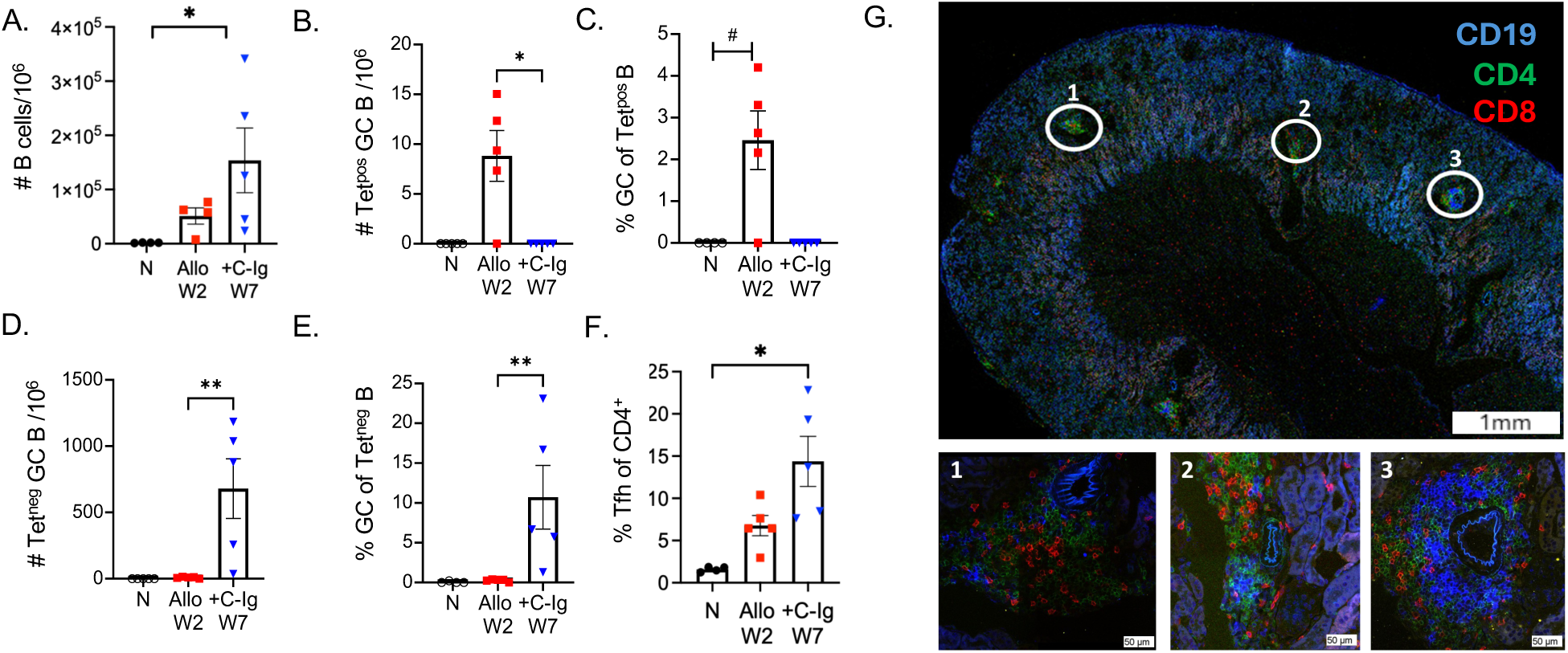
B & Tfh cells accumulate in acute rejecting (Allo) or CTLA-4Ig-treated (+CTLA-4Ig) B/c kidney allografts. (A) Number of graft-infiltrating B cells recovered normalized to 10^6^ recovered lymphocytes. (B) Number of graft-infiltrating tetramer positive (Tet^pos^) GC B cells recovered normalized to 10^6^ recovered lymphocytes. (C) Percentage of GC (Fas^+^GL7^+^) of donor-MHC Tet^pos^ B cells. (D) Percentage donor-MHC Tet^neg^ of total B cells. (E) Number of graft-infiltrating Tet^neg^ GC B cells recovered normalized to 10^6^ recovered lymphocytes. (F) Percentage of Tfh (CXCR5^+^PD1^+^) cells of total CD4^+^ T cells. Statistical significance was assessed with 2-way ANOVA with Dunnett’s multiple comparison test. *P < 0.05; ****P < 0.0001 or Student’s t-test ^#^P < 0.05.

### Intrarenal autoreactive B cells sense antigen locally in the graft

To assess whether intrarenal B cells were encountering antigen within the graft, we utilized Nur77^GFP^ reporter mice wherein we first confirmed that *in vitro* BCR engagement with anti-IgM induced transient endogenous Nur77 and GFP expression (Fig. 5A-B)^44,45^. Furthermore, Nur77 and GFP were more rapid readouts of BCR engagement compared to CD69 expression (Fig. 5C), and the GFP reporter was slightly more sensitive than endogenous Nur77. Importantly, the levels of both Nur77 and GFP trended downwards at 24h consistent with transient induction by recent BCR engagement (Fig. 5D).

**Figure 5.**
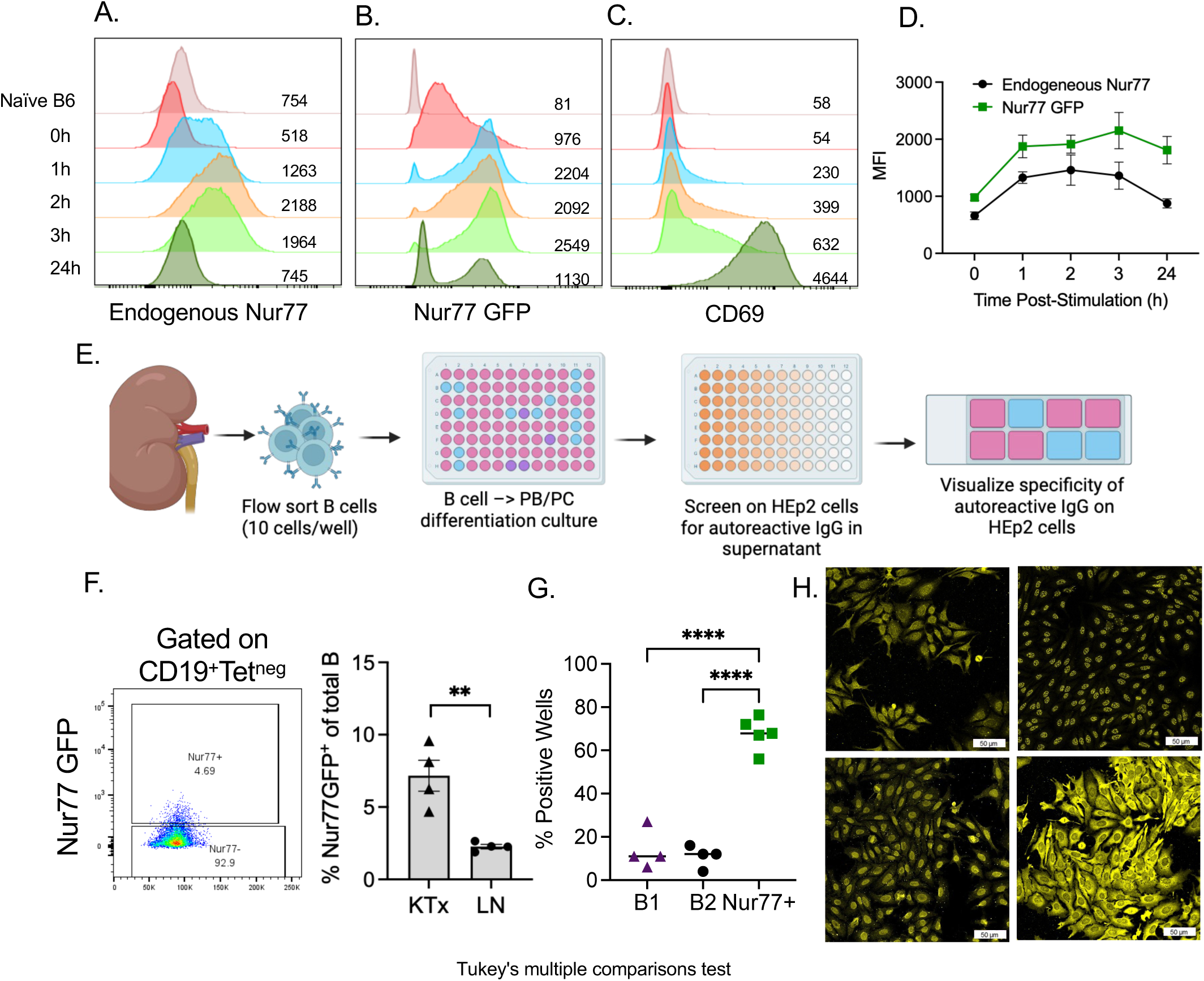
Autoreactive B cells sense antigen and upregulate Nur77 in BALB/c kidney allografts. (A-C) Representation of Endogenous Nur77, GFP and CD69 expression in B cells from Nur77^GFP^ transgenic mice after stimulation with anti-IgM F(ab’)_2_ + anti-CD40 for the indicated times (Y axis). Numbers within panels indicate MFI. (D) Quantification of endogenous Nur77 and GFP expression by CD19^+^ B cells. (E) Schematic of flow sorted Nur77-GFP^+^ CD19^+^ B cells from CTLA-4Ig-treated recipients were placed in Nojima cultures (10 cells/well), and autoantibody detection in culture supernatants using HEp-2 ELISA. (F) Gating strategy of flow sorted Nur77^GFP+^ B cells, and percentage of Nur77^GFP+^ of total intrarenal and LN B cells. (G) Percentage wells with autoreactive IgG in culture supernatants of intrarenal B1 (B220^+^CD11b^+^), B2 (B220^+^CD11b^neg^) or Nur77^GFP+^ CD19^+^ B cells. (H) Representative confocal images of autoreactive IgG from culture supernatants binding to HEp-2 cells. Statistical significance was assessed by ordinary ANOVA test and Tukey’s multiple comparison test. **P < 0.01; ****P < 0.0001.

We next interrogated the expression of Nur77^GFP^ by intrarenal or lymph node B cells isolated from CTLA-4Ig-treated BALB/c kidney recipients at 8 weeks post-transplant (Fig. 5E). Approximately 5-10% of CD19^+^ B cells from rejecting kidney grafts had elevated Nur77^GFP^ expression and were predominantly MHC tetramer-negative B cells (Fig. 5F, Fig. S6A-B), consistent with preferential antigen sensing by non-donor MHC-specific B cells. Finally, a significantly lower percentage of lymph node CD19^+^ B cells demonstrated elevated Nur77^GFP^ expression compared to intrarenal CD19^+^ B cells (Fig. 5F).

To confirm that intrarenal Nur77^GFP^⁺ B cells were enriched for autoreactivity, flow-sorted Nur77^GFP^⁺ CD19^+^ B cells were placed into Nojima cultures (10 cells/well) that allow B cells to differentiate into antibody-secreting cells ^46–48^. After ∼10-12 days in culture, the supernatants were tested for HEp-2 reactivity (Fig. 5E). Because transcriptional analysis by Asano et al. ^13^ indicated that intrarenal B cells were enriched for an innate-like phenotype, we also included controls of Nur77^GFP–^ B cells that were subdivided into B1 vs B2 cells (Fig. S6C) based on B220 and CD11b expression ^49^. Approximately 70% of wells with Nur77^GFP+^ B cells that were confirmed to have IgG in the supernatant also had HEp-2–reactive IgG; in contrast Nur77^GFP^⁻ B1 or B2 cells were rarely HEp-2-positive (Fig. 5G). HEp-2 positivity was specific for cytoplasmic, nuclear or nucleolar antigens, or were broadly reactive (Fig. 5H), and reflected the range of reactivities observed with sera of mice undergoing transplant rejection. Together, these findings show that autoreactive B cells within rejecting grafts engage autoantigen to differentiate to autoantibody-secreting cells.

### IL-15 supports intrarenal B cell accumulation and autoantibody production

We addressed the possibility that autoantibodies generated during rejection could have direct pathogenic activity. We first tested whether sera from rejecting and CTLA-4Ig–treated recipients with high HEp-2 reactivity (binding to intracellular antigens; Fig. 1H-I) could also bind to surface-accessible antigens expressed by live, non-permeabilized HEp-2 cells using the more sensitive flow cytometry assay (Fig. S7A-B). We observed a modest increase in auto-reactive IgG binding to live HEp-2 cells in sera from rejecting recipients compared to sera from naive mice (Fig. S7C-D). We next tested if autoreactive IgG from the sera of CTLA-4Ig-treated recipients or naive mice. were capable of inducing complement-mediated cytotoxicity. To eliminate the potential contribution of DSA, we used primary renal endothelial cells from recipient C57BL/6 mice as targets. Incubation of endothelial cells with sera cells was followed by incubation with rabbit complement. A significant increase in cell death was observed with sera from CTLA-4Ig-treated kidney transplant recipients compared to naïve sera (Fig. S7E-F). These results suggest that graft-derived autoantibodies could mediate complement-dependent injury of renal endothelial cells, supporting a potential direct, albeit modest, pathogenic role for autoreactive IgG generated during kidney allograft rejection. These observations raise the possibility that preventing autoantibody production could have salutary effects on the transplanted kidney graft.

Asano et al. ^13^ reported that class-switched intrarenal B cells expressed the innate pro-inflammatory cytokine IL-15, and that IL-15 was readily detected by immunofluorescence in infiltrating B and other immune cells in rejecting human renal allograft biopsies. Moreover, neutralization of IL-15 has been reported to limit kidney transplant rejection ^17^. Given the potential pro-inflammatory role for IL-15, we tested whether early blockade of IL-15 could reduce intra-graft autoantibody production and improve graft outcomes. Some CTLA-4Ig–treated recipients were administered an IL-15–blocking monoclonal antibody^50^ on post-transplant days 0, 3 and 7, and intragraft immune cells as well as autoantibody production were assessed at 6-8 weeks post-transplant. Anti-IL-15 in combination with CTLA-4Ig significantly improved the quality of the kidney grafts and reduced the incidence of hydronephrosis, i.e. swelling of kidney due to urine buildup subsequent to blockage in the urinary tract, compared to CTLA-4Ig alone (Fig. 6B-C). Anti-IL-15 also substantially reduced infiltrating B220^+^ B cells as well as CD4^+^ and CD8^+^ T cells (Fig. 6D-E). Consistent with overall reduced cellular infiltrates, kidney organoids from anti-IL-15-treated recipients produced significantly less autoreactive IgG (Fig. 6F-G). Together, these data implicate IL-15 as a supportive factor for local autoantibody production and the intrarenal accumulation of T and B cells in rejecting kidney allografts.

**Figure 6.**
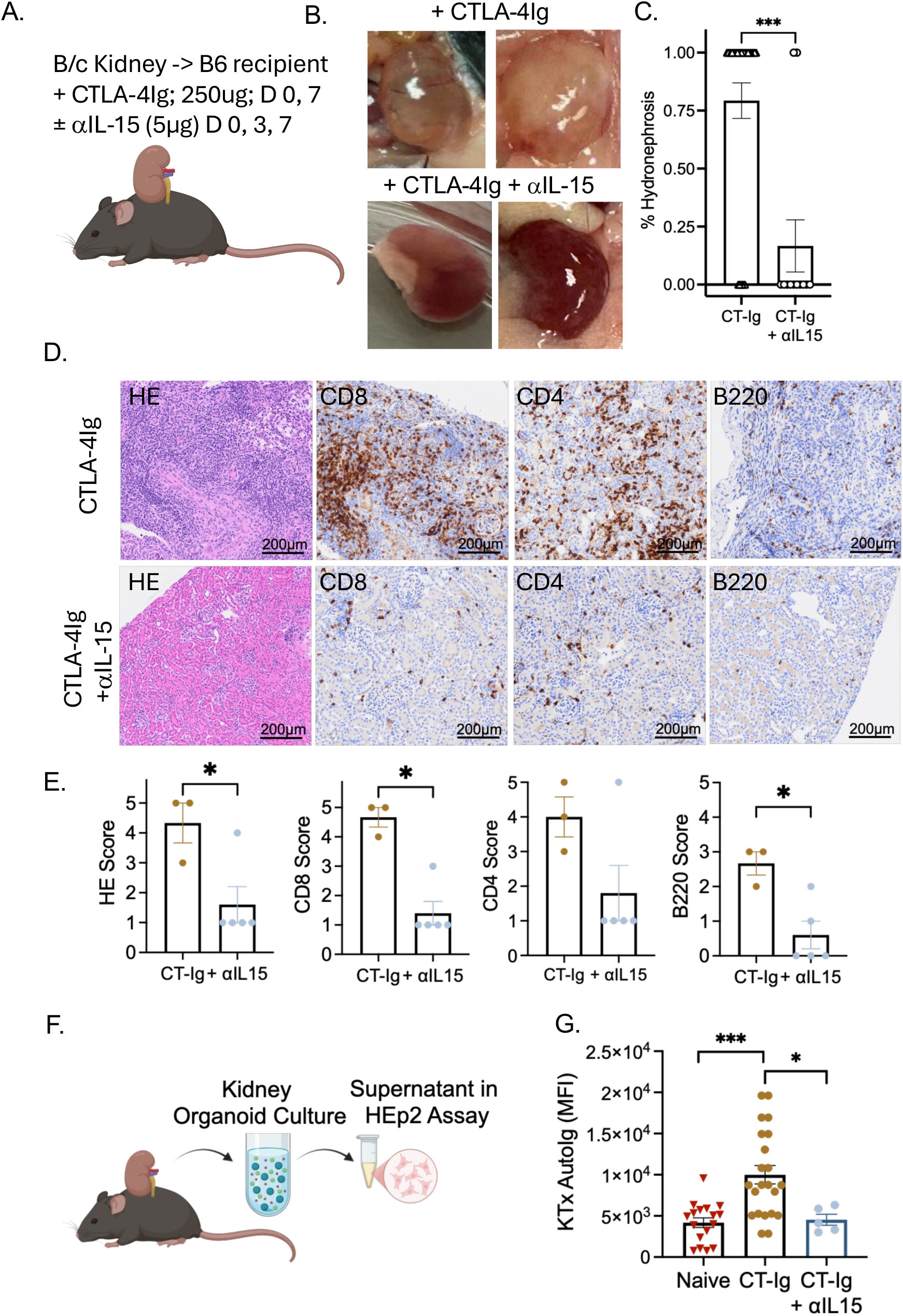
Transient anti-IL15 and CTLA-4Ig induces long-term suppression of intra-kidney autoreactive IgG production. (A) Experimental design. (B) Representative whole kidney images and (C) Kidneys with hydronephrosis (1) or no hydronephrosis (0). Each symbol represents one recipient (n=29 for CTLA-Ig (CT-Ig); n=12 (CT-Ig + anti-IL15 (aIL15). (D) Representative images or quantification of HE or immunohistochemistry staining for CD8, CD4 and B220 (0=minimal infiltration; 4=maximum infiltration). (F-G) Autoreactive IgG (AutoIg) produced by kidney allograft organoids from CTLA-4Ig, or CTLA-4 Ig + aIL15 treated recipients. Naïve kidneys were from B/c donors. Each symbol represents one organoid culture, and with 2-5 organoids per recipient. Statistical significance was assessed by Mann-Whitney test. *P < 0.05; ***P < 0.001.

## Discussion

Studies by Asano et al.^13^ revealed a breach of B cell tolerance during kidney allograft rejection that led to the differentiation of intrarenal B cells into cells producing autoreactive antibodies that bound to renal-specific or inflammation-associated antigens. Furthermore, those class-switched B cells had an innate cell transcriptional state resembling mouse peritoneal B1 or B-innate cells. Those findings were congruent with emerging literature for a complementary role of non-HLA, in addition to donor HLA antibodies, in transplant rejection ^26,28,29,51–54^. We hypothesized that infiltrating innate-like B cells contributed to local tissue destruction and pathological features of AMR through the local production of pathogenic autoantibodies or by serving as antigen-presenting cells to promote T cell-mediated rejection (TCMR). This study demonstrates the unique immunobiology of autoantibody versus DSA responses in mouse renal allograft transplantation, and their potential roles in promoting allograft rejection.

Conceptually, targets for non-HLA antibodies can be divided into those that recognize polymorphic antigens that are mismatched between donor and recipient, autoantigens that undergo post-translational modifications, self-antigens that are secreted or expressed on the cell surface under inflammatory states, and ubiquitously expressed self-antigens ^22,26,55,56^. While the first 2 classes of non-HLA antibodies are by definition non-self-antigens, most assays lack the resolution to distinguish minor polymorphic differences or post-translational modifications on self-antigens. Recent studies by Zorn and colleagues ^57,58^ have characterized some of these modifications, wherein monoclonal antibodies generated from neonatal B cells were shown to recognize common chemical adducts exposed on modified amino acids and other macromolecules. In addition, proteoform imaging mass spectrometry are revealing post-translational modifications that are tissue specific ^55,56^. Finally, earlier findings that intrarenal B cells recognize Ki-67 and HEATR antigens that are induced in inflamed tissues are supported by recent findings that intragraft plasma cells recognize bilirubin, a heme-degradation byproduct that accumulated in smooth muscle cells of transplanted hearts with vasculopathy but was not expressed in healthy heart tissue^24^.

The notion that the number of non-HLA antigens is likely to be much larger than HLA antigens is supported by proteomic array analysis by Sarwall and colleagues^59^, who also reported that a subset of non-HLA antibodies recognizing antigens in the renal cortex was significantly associated with the development of chronic kidney allograft injury^60^. Moreover, in murine kidney allografts, less than half of intrarenal switched B cells were alloreactive, consistent with the remaining B cells being specific for non-HLA antigens and/or were non-specifically accumulating in inflamed grafts ^17^. The broad array of non-HLA antigens capable of eliciting antibody responses in the transplant setting evokes two related but currently unresolved questions: which is the best assay for detecting non-HLA IgG, and what specificities of non-HLA antibodies are most pathogenic. In our study, we assessed autoantibodies using a commercial anti-HEp-2 ANA kit, which allowed us to test antibody binding to a broad range of intracellular antigens. We further modified this assay from a labor-intensive confocal microscopy approach to a flow cytometry-based assay that was more tractable and quantitative, thereby allowing serum dilutions to be performed to more rigorously assess autoreactive IgG titers ^61^. We also addressed the concern that HEp-2 cells are HeLa human cervical cancer cells by also showing that sera binding to HEp-2 cells also bound to autoantigens expressed by mouse kidney cells. We did not observe any false negatives (binding the HEp-2 but not to kidney tissue), however, approximately 30% of sera were autoreactive only to mouse kidney cells and were missed by the HEp-2 assay, indicative of autoreactive IgG that recognized kidney-specific targets. Finally, we showed that the majority of sera that bound to HEp-2 cells were negative in the non-HLA bead assay which assesses only 60 target antigens. The concern of interspecies reactivity which is partly addressed by the HEp-2 also being of human origin, and mouse kidney studies discussed above. The utility of the non-HLA bead assay was its ability to show increasing IgG MFI to specific autoantigens and epitope spreading with time post-transplantation. Thus, the HEp-2 assay detects a broader repertoire of non-HLA autoantibodies, while the bead assay captures the evolution of the autoreactive antibody repertoire post-transplant.

Intrarenal accumulation of autoreactive B cells suggested that autoreactive antibodies could be produced within rejecting kidneys regardless of whether serum DSAs were detected ^13^. Indeed, our studies in mouse kidney transplant models showed that DSA and autoantibody production were independently regulated. Furthermore, while DSA and autoantibodies were produced in acute kidney allograft rejection, autoantibodies were produced even when DSA production was inhibited by transient CTLA-4Ig. We confirmed these findings with a single MHC Class II-mismatch kidney transplant model involving bm12 kidneys in C57BL/6 recipients, thus extending the observations of Pettigrew and colleagues of autoantibody production in a bm12 into C57BL/6 heart transplant model ^32^. We also showed that autoreactive antibodies were produced preferentially within rejecting kidneys, whereas DSAs were produced in both draining lymph nodes and kidney grafts, especially early post-transplant. Importantly, the production of autoantibodies within the kidney persisted with time post-transplantation in CTLA-4Ig-treated recipients. Finally, we provide evidence for the accumulation of autoreactive B cells in the kidney, and for antigen recognition that allowed for B cell receptor directed upregulation of Nur77. Indeed, co-occurrence of B cells with germinal center phenotype (Fas^+^GL7^+^), T cells with Tfh phenotype (CXCR5^+^PD1^+^) and lymphoid aggregates suggests potential sites facilitating autoantibody production within rejecting kidney allografts. These observations complement observations of IL-21-producing effector Tfh cells and transcriptionally distinct B cells accumulating within acutely rejecting kidney allografts and contributing to *in situ* DSA responses ^16, 48^.

Class-switched B cells that accumulated in human renal allografts expressed IL-15 transcripts, and immunohistochemical staining confirmed that intrarenal B cells but not tonsil germinal center B cells expressed IL-15 and that renal tubular cells expressed IL-15R ^13^. These observations suggested a model whereby IL-15 secreted by B cells under inflammatory conditions were captured by tubular cells for presentation to immune cells, thus providing an additional mechanism of graft damage ^62–64^. This model raised the possibility that neutralization with anti-IL-15 might prevent autoantibody production and preserve kidney allograft pathology ^63,65,66^. Indeed, administration of a neutralizing anti-IL-15 together with CTLA-4Ig in the first week post-transplantation prevented local autoantibody production and T cell infiltration, and significantly improved kidney graft pathology. These observations extend the findings of Tieu et al. ^63^ that tissue-resident memory T cell maintenance in kidney allografts requires both cognate antigen and IL-15. Finally, while transcriptionally and functionally distinct and/or innate-like B1 cells accumulate in rejecting kidneys ^13,48^, our studies showed that B1 cells were not enriched for autoreactivity. Whether B1 cells express anti-inflammatory molecules such as IL-10, PDL1 and CTLA-4, and play immunoregulatory roles, as recently reported by Shen et al. ^67^ requires further investigation.

A major question confronting studies on autoantibodies in allograft transplantation is whether they are pathogenic, or if their production is a sentinel of intrarenal inflammation and cell damage. Our studies suggest that autoantibodies can have a pathogenic role by binding to cell surface autoantigens and mediating antibody-mediated cytotoxicity, similar to MHC-specific antibodies. However, the vast majority of autoantibodies bind to nuclear and cytoplasmic targets, leads us to speculate that they primarily bind to dying cells to generate opsonins that facilitate antigen uptake and presentation by antigen-presenting cells to graft-specific T cells. Indeed, the observation that anti-IL15 + CTLA-4Ig treated recipients lack T cell infiltrates observed in CTLA-4Ig treated recipients supports this possibility. Nevertheless, definitive studies are necessary in both mouse models and human organoid cultures to clarify the function of autoantibodies in allograft rejection, and the roles played by the local production of IL-15 by intrarenal B cells.

In summary, our studies build on the observational findings with human kidney biopsies ^13^ to support a mechanistic model whereby local inflammation prompts a breach in organ-restricted B cells tolerance, resulting in the production of autoantibodies that then promote local tissue destruction. Several features of kidney allograft rejection could plausibly foster this loss of B cell tolerance and production of autoantibodies, including ischemia-reperfusion injury, persistent alloantigen presentation, exposure to damage-associated molecular patterns, and local production of inflammatory cytokines such as IL-15. These features might be replicated in other disease settings with chronic inflammation and tissue damage. Our studies also underscore differences in immunobiology of DSA versus autoantibody production and the unique technical challenges of capturing their broad specificity. Finally, that autoantibodies are produced in the kidney allograft suggests that their detection in the serum substantially underestimates the concentration and range of antigen specificity present within the allograft where they are likely to have the most potent effects. These limitations handicap studies attempting to correlate autoantibody production with graft outcomes and underscore the need for accurate assessments of *in situ* autoantibody production in clinical transplantation.

## Material and Methods

### Sex as a biological variable

Our study examined only female animals because kidney transplants could only be successfully performed in female mice. It is unknown whether the findings are relevant for male mice.

### Mice

Eight- to 12-wk-old female C57BL/6 (B6, H-2b) mice and 6- to 8-wk-old female BALB/c (B/c, H-2d) mice were purchased from the Harlan Laboratories. Nur77-GFP (B6N.B6-Tg(Nr4a1-EGFP/cre)820Khog/J) mice ^45^ and bm12 (B6(C)-H2-Ab1bm12/KhEgJ) mice were purchased from Jackson Laboratories. CD19^Cre^ and Blimp-1^flox/flox^ mice were purchased from Jackson Laboratories and bred inhouse ^33,68^.

### Kidney transplantation

Kidney transplantation was performed with kidneys from BALB/c mice transplanted into C57BL/6 (H-2^b^) recipients that had been subjected to left nephrectomy. In this model, the recipient retains one functioning kidney graft for the entirety of the experiment to ensure the mice do not develop uremia when the allografts were rejecting, since uremic states can result in immune dysfunction ^69,70^. On day 0 and day 7 post-transplant, CTLA-4Ig (Abatacept, Bristol-Myers Squibb) was given by intraperitoneal injection (250 µg/mouse), while neutralizing anti-IL-15 (Clone AIO.3, BE0315; BioXCell ^50^) was administered on day 0, 3 & 7 by intraperitoneal injection (5 µg/mouse). All animals were housed in pathogen-free conditions in accordance with the University of Chicago Institutional Animal Care and Use Committee.

### DSA quantification

Sera from naive or rejecting mice were screened for donor-specificity using BALB/c splenocytes as targets and results are presented as mean fluorescence intensity (MFI) of mouse IgG binding to non-B cell targets. We also used a multiplex single antigen bead assay to quantify reactivity to donor MHC Class I and Class II as previously reported ^30,42^. In brief, streptavidin-coated multiplex beads (PAK-5067-10K; Spherotech) were tetramerized with biotinylated H-2K^d^, H-2L^d^, I-E^d^, I-A^d^ or control I-A^b^ monomers (NIH tetramer core facility). Sera or culture supernatants were incubated with MHC-coated beads and then stained with FITC-conjugated anti-mouse IgG (406001; Biolegend). The MFI of IgG binding was determined by flow cytometry (Novocyte Quanteon, Agilent)

### Immunofluorescence Staining and Imaging

Autoantibodies were detected using the HEp-2 ANA detection kit (26103, BIO-RAD Laboratories). Briefly, HEp-2 cells were incubated with patient serum and rabbit anti-human ENO1 (11204-1-AP, Proteintech). After washing, cells were incubated with AF568 goat anti-mouse IgG (A11004, Invitrogen), Hoechst 33342 (62249, Thermo Fisher Scientific), and AF647 donkey anti-rabbit IgG (A32795, Invitrogen). Slides were imaged using the Stellaris 8 Falcon confocal microscope under identical acquisition settings across all samples.

### HEp-2 Cell Image Analysis Using CellProfiler

A CellProfiler colocalization pipeline was used to analyze the regions of interest (ROI) and quantify relative anti-mouse IgG binding intensities (https://cellprofiler.org/published-pipelines). This pipeline comprised of three channels, the first being Hoechst 3342 to stain the nucleus while the second was anti-Enol-1 which stained the cytoplasm. These two channels played an integral part of cell segmentation process. The third channel was the autoantibody channel, which was overlayed with the nuclear and cytoplasmic channels to assess the areas of the cell that the autoantibodies were targeting. In this pipeline, the IdentifyPrimaryObject module was used to find the nuclear area and the IdentifySecondaryObject module was used for the cytoplasmic area. The method used for the secondary object was through “propagation” and the threshold strategy was “adaptive” with an “Otsu” thresholding method. A Hoechst channel grayscale image was used to identify the nuclear primary object, and an ENO-1 grayscale image was used to identify the secondary object. The OverlayObjects module was used to overlay the autoantibody channel with the initial two channels for the identification of intensity and location of autoreactive binding. The quantitative data was then exported to Excel via CellProfiler, and data were plotted using GraphPad Prism.

### Tissue Image Analysis Using Custom Python Workflow

Kidney tissue images were processed using a custom Python-based image analysis pipeline to quantify nuclear and cytoplasmic fluorescence intensities. The nuclear channel (_ch00.tif), with Hoechst staining, was smoothed using a Gaussian filter to reduce noise and thresholded to generate a binary nuclear mask. A second Gaussian-filtered image was used to define a broader tissue mask. The cytoplasmic compartment was computationally defined by subtracting the nuclear mask from the tissue mask, thereby isolating non-nuclear tissue regions.

To correct for uneven illumination and background fluorescence in the autoantibody channel (_ch01.tif), a large-scale Gaussian blur was applied to estimate background intensity. The original autoantibody image was normalized to this blurred image and rescaled to the 16-bit dynamic range. Total fluorescence intensity within nuclear and cytoplasmic regions was calculated by applying the respective masks to the corrected image. Pixel counts were used to determine compartment areas, enabling calculation of mean nuclear and cytoplasmic intensities. The nuclear-to-cytoplasmic mean intensity ratio was then computed to assess relative autoantibody localization within tissue sections.

Both the HEp-2 CellProfiler pipeline and custom Python tissue-analysis code are publicly available at:https://github.com/Deepgh/Autoantibody-quantification.

### Autoantibody detection with flow cytometry assay

Cultured HEp-2 cells were detached from plates by 0.05% trypsin or Accutase (BDBiosciences), stained with Live Dead Aqua (Invitrogen) then fixed and permeabilized by Fix/Perm buffer set according to the manufacturer’s instruction (eBioscience 00-5123-43). Detached cells (100,000 – 200,000 per sample) were incubated with serially diluted mouse sera and rabbit-anti human ENO-1 (11204-1-AP, Proteintech) and washed. Cross-adsorbed PE goat anti-mouse IgG (P-852, Invitrogen) and AF647 donkey anti-rabbit IgG (A32795, Invitrogen) were used as detection antibodies. Flow cytometry analysis was performed on the Attune Penteon/Quanteon analyzer.

### Autoantibody detection with Lifecode non-HLA antibody assay

The detection of autoantibodies with LIFECODES Non-HLA Antibody Kit (Werfen), which covers 60 target antigens, following manufacturer’s recommended protocols with the exception that anti-mouse IgG PE (Invitrogen, P-852) was used as detection antibody.

### Indirect immunofluorescence on frozen mouse kidney tissue sections

The left kidney of CD19^Cre^Blimp-1^flox/flox^ mice (JAX) were subjected to warm ischemia reperfusion injury (IRI) for 30 minutes by clamping the renal artery. Mice were sacrificed at 30 days later and both IRI and non-IRI kidneys were harvested. Kidneys were fixed in 1%PFA overnight and transferred into 30% sucrose solution. Kidneys were then embedded in OCT and sectioned into 5 μm thickness and rehydrated in PBS. Each section was then blocked overnight using TruStain FcX PLUS (anti-mouse CD16/32; 156604, Biolegend) in 0.1% Triton-X (9002-93-1, Sigma), 1% BSA (9048-46-8, Sigma), then incubated with serum. After washing, slides were stained with Hoechst; 33342 (62249, ThermoFisher Scientific) and AF568 goat anti-mouse IgG (A11004, Invitrogen).

Transplanted kidneys were stained with a combination of primary antibodies: rat anti-CD19 (115538, 1:100, Biolegend), rat anti-CD4 (100423, 1:100, Biolegend), rat anti-CD8b (126612, 1:100, Biolegend). Stained sections were mounted in ProLong Gold Antifade Mountant (P36934, ThermoFisher Scientific) and imaged with Stellaris confocal microscopy (Leica).

### Flow cytometry of intrarenal T and B cells

Fresh kidney grafts were dissociated by enzymatic dissociation using collagenase IV and DNAse I (9001-1201, 9003-98-9, Sigma-Aldrich). The following antibodies were used for surface staining at 4°C for 30 minutes: Biotinylated H-2K^b^ monomers, or H-2K^d^, I-E^d^, I-A^d^ and H-2L^d^ tetramers (NIH Tetramer Core Facility). Live Dead Aqua (Invitrogen), anti-CD19 (30-F11, Biolegend), anti-B220 (RA3-6B2, Biolegend), anti-IgD (11-26c.2a, Biolegend), GL-7 (GL7, Biolegend) and CD95/Fas for germinal center B cells (Jo2, BD Biosciences), anti-CD90.2 (S20008D, Biolegend), anti-CD4 (GK1.5, Biolegend), CXCR5 (L138D7, Biolegend) and PD-1 (29F.1A12, Biolegend). Samples were intracellularly stained with anti-Bcl-6 (K112-91, eBiosciences) after fixation with the eBioscience Foxp3 Fix/Perm buffer set according to manufacturer’s instructions (00-5523-00, ThermoFisher Scientific). Flow cytometry analysis was performed on a Attune Pentaon analyzer and analyzed with FlowJo v10 (FlowJo LLC).

### Organoculture to assess in Vitro Production of DSA-and autoantibodies

In vitro organo-culture to capture production of DSA and autoantibodies were detected as previously described ^37^. Briefly, the lumbar LN and native and transplanted kidneys were harvested at the indicated times post-transplant. The renal cortex of fresh native kidney and allografts were dissected and cut with a sterile scalpel into ∼30mg fragments. A total of 2-5 tissue fragments were collected and washed three times in RPMI-1640 medium (Cambrex, Walkersvile, MD) and placed in 24-well plate in 1ml of RPMI-1640 (61870036, ThermoFisher) containing 10 % fetal bovine serum (10100147, Gibco^TM^), and 100 U/ml penicillin-streptomycin (15140122, Gibco^TM^). After 4 days incubation, the supernatant was harvested, centrifuged at 2000 r.p.m for 10 min and stored at −20℃ until antibody analysis was performed. Bortezomib (VELCADE®, Takeda) at 50nM was added at the beginning of the organoid cultures for 24 or 96 h.

### Nojima B cell cultures

Sorted Nur77^GFP+^ and Nur77^GFP–^ intrarenal B cells were cultured with NB-21.2D9 feeder cells^46,47^ (generous gift from Garnett H. Kelsoe, Duke University). NB21.2D9 cells were seeded into 96-well plates at 2,000 cells/well in B cell media: RPMI-1640 supplemented with 10% FBS (SH30070.03, Thermo), 5.5×10^-5^ M 2-mercaptoethanol (21985, invitrogen), 10 mM HEPES (15630-080, Invitrogen), 1 mM sodium pyruvate (11360-070, Invitrogen), 100 U/mL penicillin-streptomycin (15140-122, Invitrogen), and MEM nonessential amino acid (11140-050, Invitrogen). The next day (day 0), 2 ng/mL recombinant mouse IL-4 (214-14-20UG Peprotech) was added, and Nur77^GFP+^ and Nur77^GFP–^intrarenal B cells were directly sorted into each well (BD ARIA Fusion). On day 2, 50% volume of culture media from the wells was removed and fresh BCM was added. On days 3 to 8, 50% of the culture media were replaced with fresh BCM. On day 10-12, culture supernatants were harvested for DSA and autoantibody quantification.

### In vitro cytotoxic assay with mouse renal endothelial cells

Autoantibody cytotoxicity assay was performed by using primary renal endothelial cells and Cytotoxicity LDH assay kit (Dojindo Labs, Rockville, MD). Renal endothelial cells were isolated from naïve 8- to 12-wk-old female C57BL/6 mice by enzymatic digestion with collagenase IV (C4-28-100mg, Sigma-Aldrich) and Roche DNAse I (2 ng/mL, Sigma-Aldrich) as previously described ^71^. Tubules were isolated via density gradient centrifugation using Percoll solution (P1644, Sigma-Aldrich) and endothelial cell isolation was performed by second round of collagenase IV digestion. Endothelial cells were cultured in 6 well plates in endothelial cell growth media (C-22110, Sigma-Aldrich) and then transferred into 96 well plates (3000 cells/well). After overnight culture, endothelial cells were washed and incubated with rabbit serum as a source of complement, and cell cytotoxicity was determined by release of lactate dehydrogenase (LDH) in culture supernatants, following manufacturer’s protocol.

## Statistical analysis

Statistical significance analyses were conducted using GraphPad Prism. Statistical differences between experimental groups were determined using Mann-Whitney U test or ANOVA, followed by Tukey’s or Dunnett’s multiple comparison test. To evaluate whether the development of antibodies across the post-transplant times differed by the study group, an interaction term between study group and time was included in the linear mixed-effects model where the P value represents the interaction term (study group × time). Correlation analysis was by simple linear regression. P values ≤0.05 were considered statistically significant.

## Study approval

All animal studies were approved by the Institutional Animal Care and Use Committee (ACUP 71236).

## Data availability

Data can be made available upon request.

## Author Contributions

IS and JCJ participated in the design of the study, conducted most of the experiments, analyzed data and generated figures. GD and MST generated the code for image analysis, JBMO assisted in the image analysis, SSD participated in some experiments, ART led the studies with the LifeCode non-HLA bead assay, and DY performed mouse kidney transplants. MRC and ASC designed the study, PTS and AJN participated in interpretation of data, and all authors edited and provided feedback on the manuscript.

## Funding support

This work was supported by NIH grants (U19 AI 082724 [MRC], R01 AI148705 [MRC and ASC], R01AI13902462 [ASC]) and Lupus Research Alliance (MRC). SSD was supported by American Heart Association post-doctoral fellowship (24POST1191785).

## Acknowledgements

MHC tetramers were provided by the NIH Tetramer Core Facility (contract HHSN272201300006C). We thank the staff of The University of Chicago Flow Cytometry Core facility (SCR_017760), Human Tissue Resource Center (SCR_019199), and Integrated Light Microscopy Core (SCR_019197) for their assistance. LifeCode non-HLA bead Antibody Kit was a gift from Werfen (Bedford, MA). We acknowledge Desiree Elaine Austin, Feinberg School of Medicine, Northwestern University, for performing the non-HLA bead antibody assessment, and Abdu Rahman Zine, University of Chicago, for assisting in the image analysis of autoantibody binding to mouse kidney tissues.

## Conflict-of-interest statement

The authors have declared that no conflict of interest exists.

## Abbreviations

AMR: antibody-mediated rejection
ANA: anti-nuclear antibody
AT1R: angiotensin II Type 1 Receptor
CTLA-4Ig: cytotoxic T-lymphocyte-associated antigen 4 immunoglobulin
DSA: donor specific antibody
GC: germinal center
HLA: human leucocyte antigen
IRI: ischemia reperfusion injury
KTx: kidney transplant
MFI: mean fluorescence intensity
MHC: major histocompatibility complex
TCMR: T cell-mediated rejection
Tfh: T follicular helper

---

**Figure S1.**
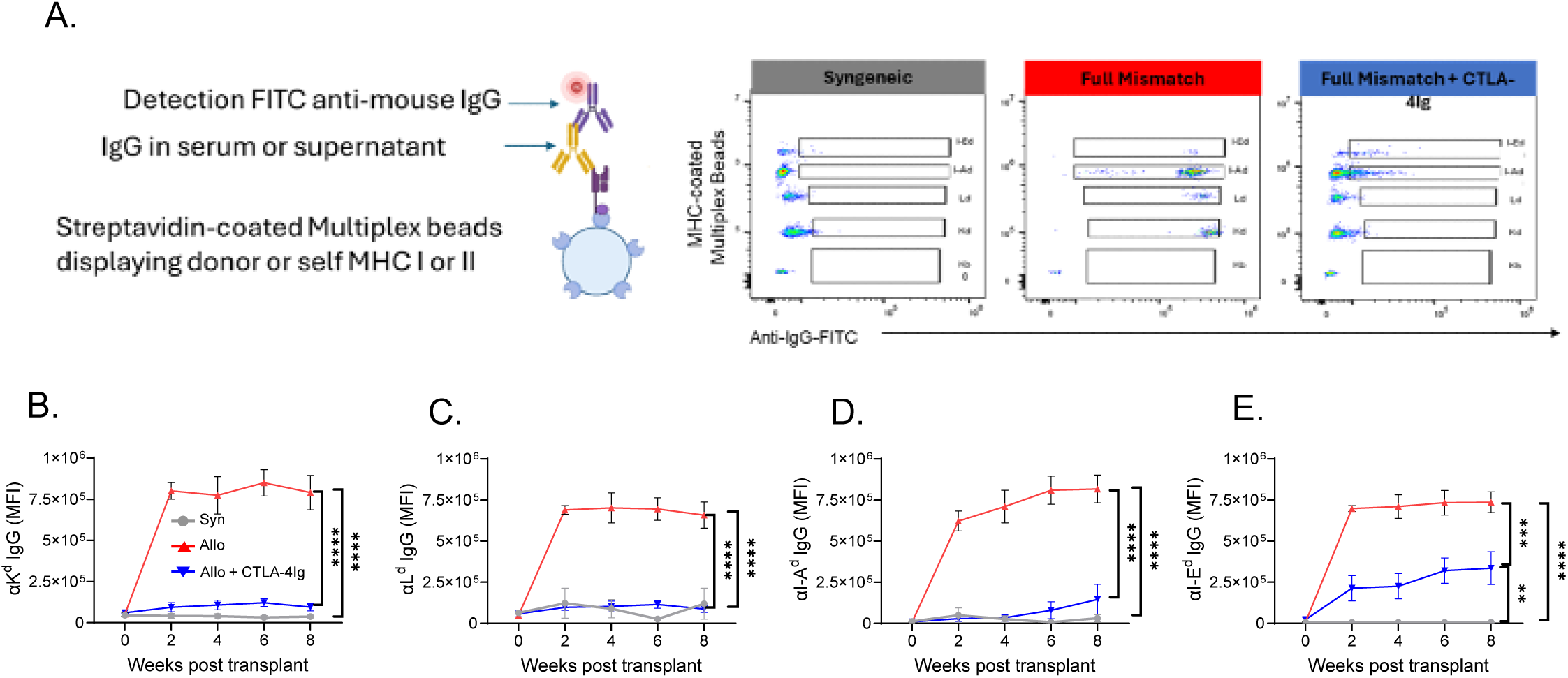
Anti-donor MHC Class I (K^d^, L^d^) and II (I-A^d^, I-E^d^) IgG production are suppressed by transient CTLA-4Ig. A. Representative flow cytometry data of anti-donor MHC specific IgG using multiplex bead assay. (B-E) Anti-donor MHC class I and Class II IgG presented as mean fluorescence intensity (MFI; n = 4 Syn or Allo + CTLA-4Ig group, n = 5 Allo). Symbols indicating experimental groups are the same for B-E. Data are presented as mean ± SEM, and statistical significance was assessed by 2-way ANOVA test. **P < 0.01; ***P < 0.001; ****P < 0.0001.

**Figure S2.**
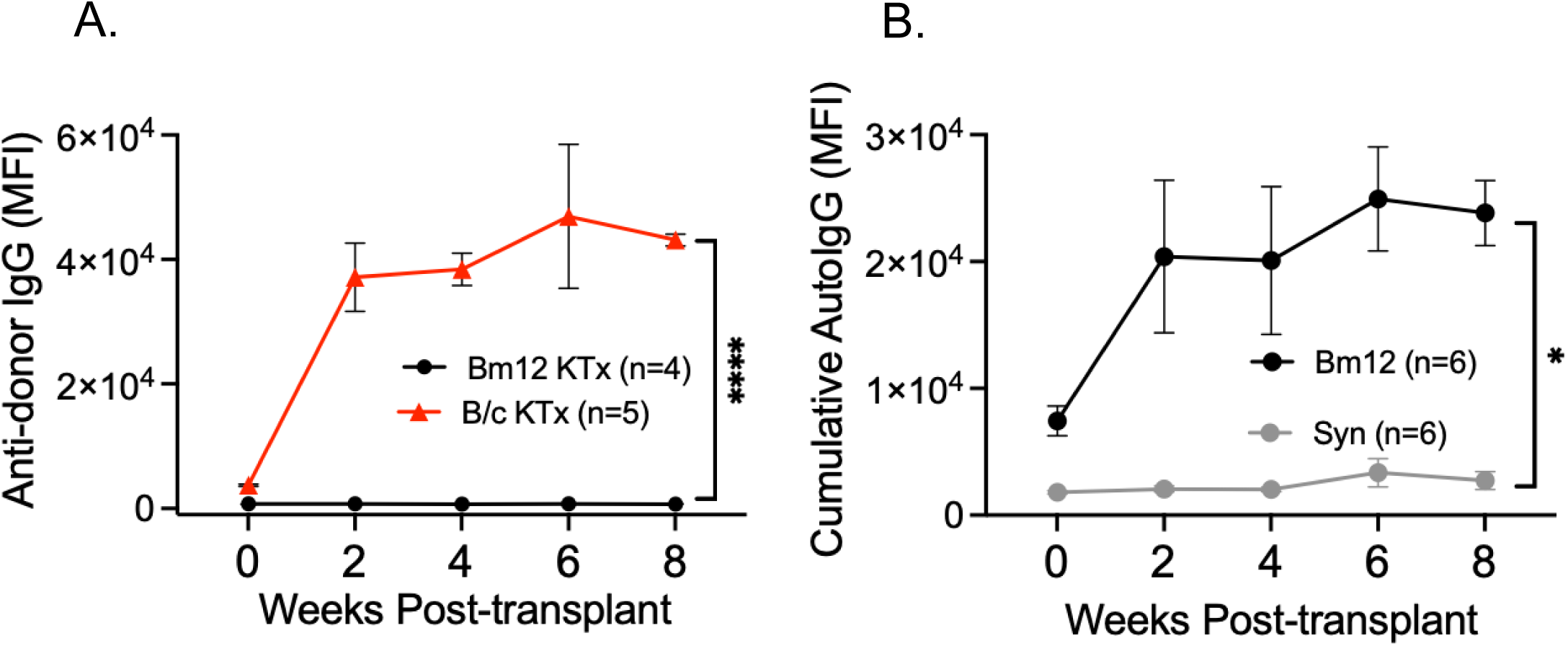
Autoreactive IgG are produced in the absence of DSA by B6 recipients of Bm12 kidney allografts. (A) Quantification of serum anti-Bm12 IgG by flow cytometry using Bm12 splenocytes as targets (n = 4). Positive control sera were from recipients of B/c kidney grafts and B/c splenocytes were used as targets (n = 5). (B) Autoreactive IgG in serum from B6 recipients of syngeneic (Syn; n = 6) or Bm12 kidney (n = 6) grafts. Data are presented as MFI ± SEM, and statistical significance was assessed by Mann-Whitney and Mixed-effects analysis. *P < 0.05; ****P < 0.0001.

**Figure S3.**
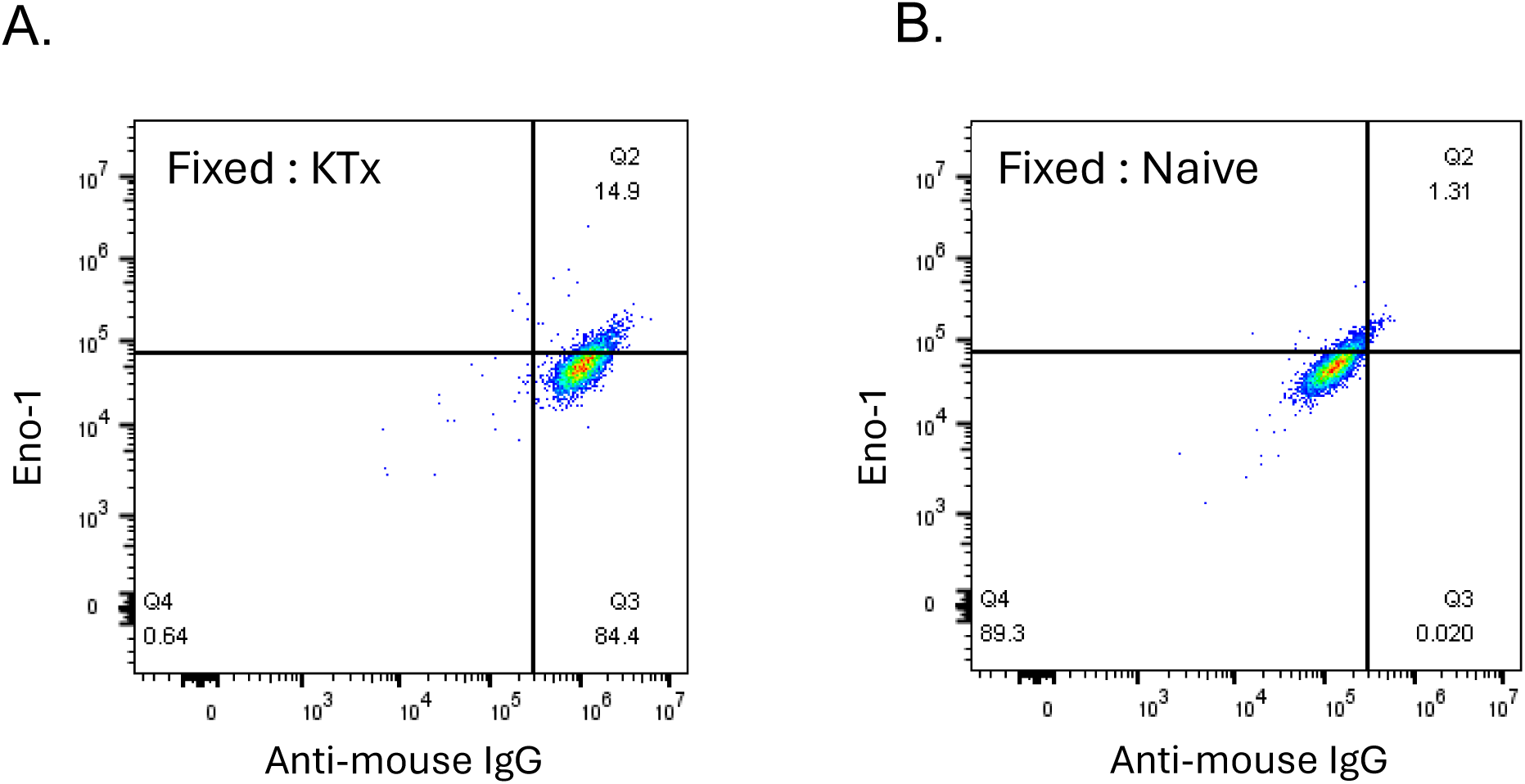
Autoantibody binding on fixed permeabilized HEp2 cells by flow cytometry. Representative flow cytometry of serum IgG (1:450 dilution in PBS) from (A) Allo + CTLA-4Ig kidney transplanted or (B) naïve mice.

**Figure S4.**
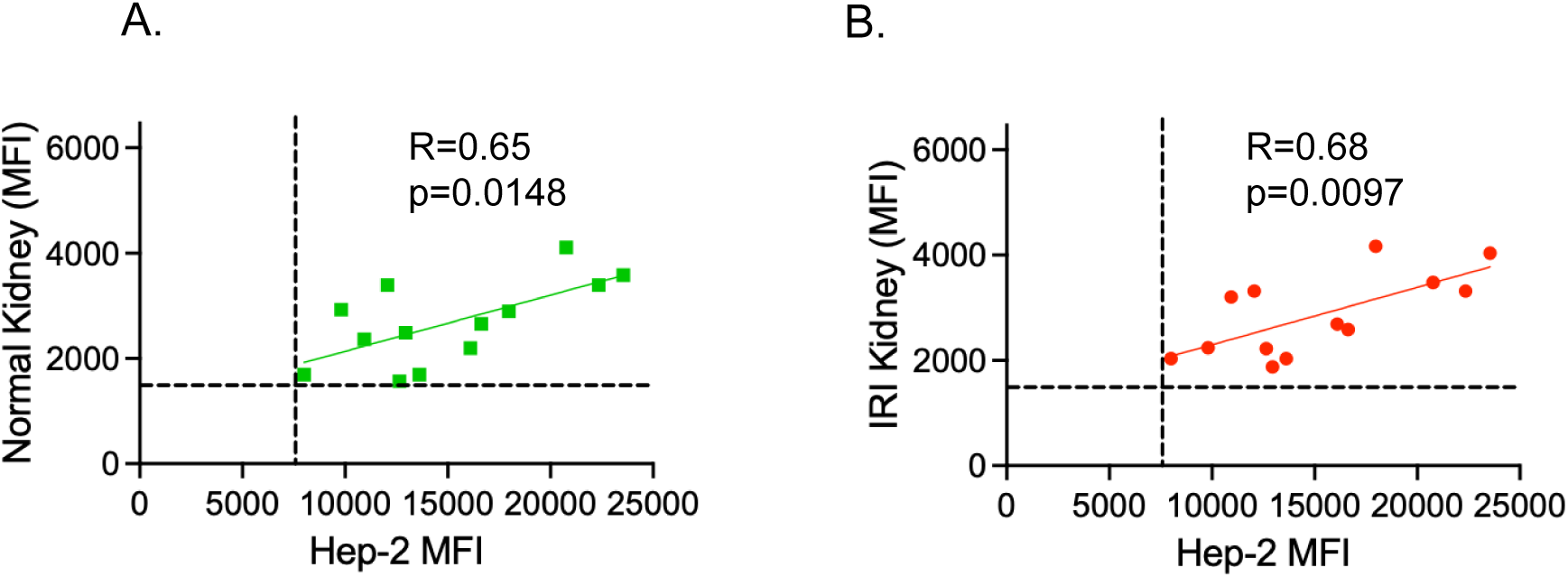
Serum HEp2-reactivity correlates with reactivity to (A) B6 normal and (B) ischemia-reperfusion injured (IRI) kidney. (n=13 matched sera). Correlation was assessed by simple linear regression.

**Figure S5.**
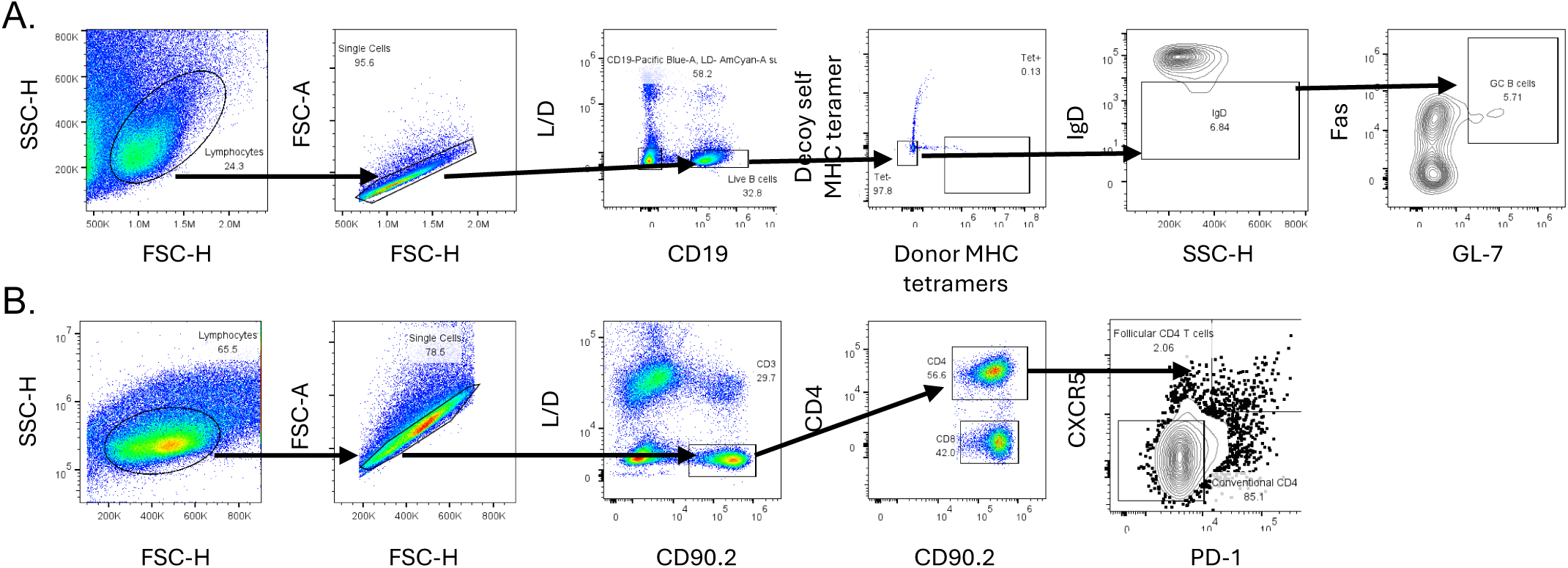
Gating strategy for graft-infiltrating (A) donor-MHC-binding B cells, and (B)Tfh cells.

**Figure S6.**
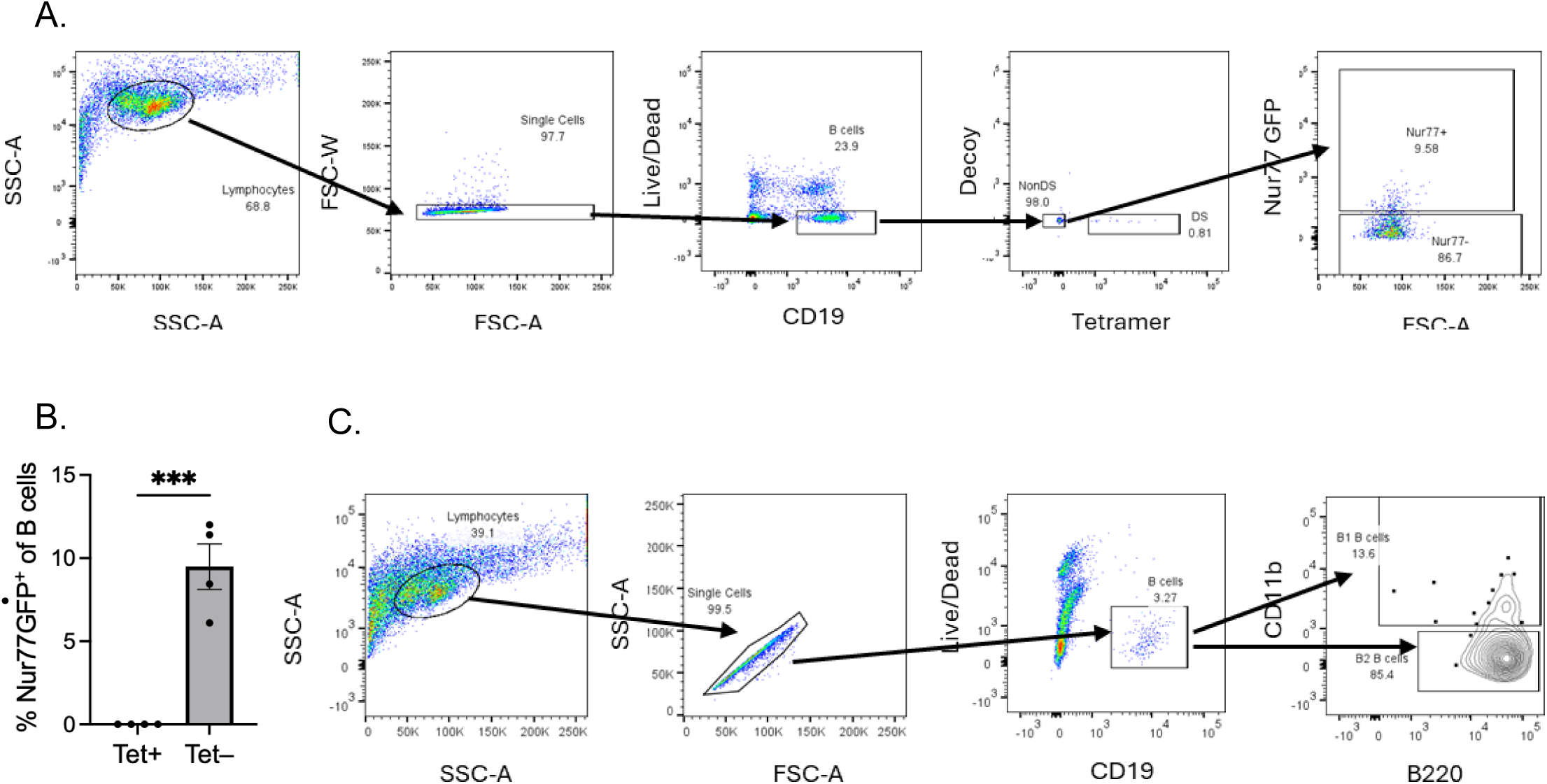
Gating strategy for graft-infiltrating (A) donor-specific B cells based on binding to pooled donor MHC (H-2K^d^, H-2L^d^, I-E^d^, I-A^d^) tetramers or non-donor-specific B cells. (B) Percentage of Nur77-GFP^+^ of tetramer-binding (Tet^+^) or negative (Tet^-^) B cells. Statistical significance determined by unpaired T test. ***P =0.0002. (C) Gating strategy for B1 (B220^+^CD11b^+^) and B2 (B220^+^CD11b^neg^) cells.

**Figure S7.**
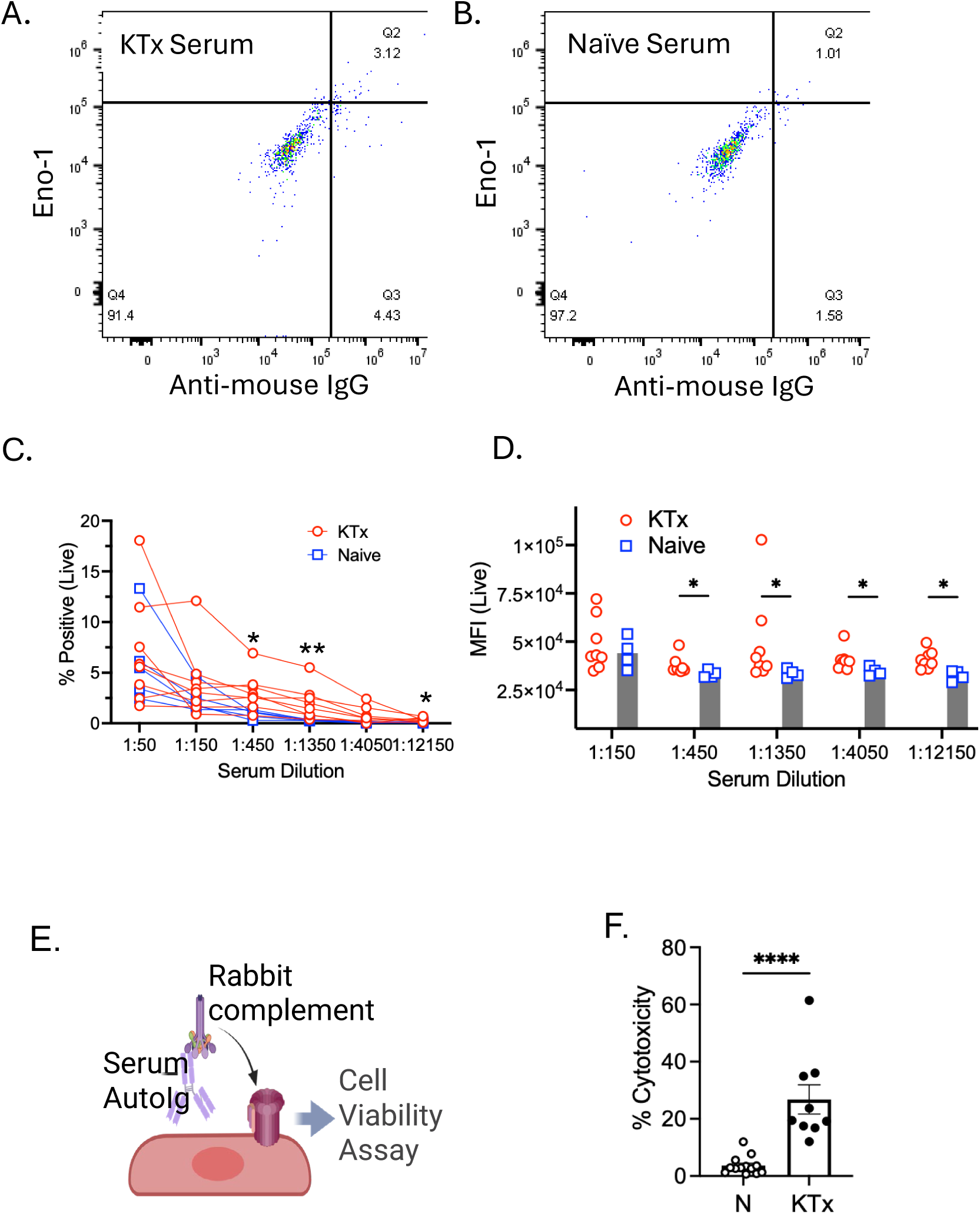
Complement-mediated lysis by autoreactive IgG. (A, B) Flow cytometric measurement of serum IgG binding to live HEp2 cells. Representative flow cytometry of serum IgG (1:450 dilution in PBS) from (A) Allo + CTLA-4Ig kidney transplanted or (B) naïve mice. Data are presented as (C) percentage positive or (D) MFI. (D). Schematic of autoreactive IgG-containing serum killing LDH assay (E) % cytotoxicity of naïve and rejecting kidney serum. Statistical significance was assessed by Mann-Whitney test. **P<0.01, *P<0.05, ****P < 0.0001,

## Notes

### Competing Interest Statement

The authors have declared no competing interest.

